# SYSTEMIC FRATAXIN DEFICIENCY CAUSES TISSUE-DEPENDENT IRON HOMEOSTASIS ALTERATIONS: IMPLICATIONS FOR FRIEDREICH ATAXIA

**DOI:** 10.1101/2024.07.05.602088

**Authors:** Maria Pazos-Gil, Marta Medina-Carbonero, Arabela Sanz-Alcázar, Marta Portillo-Carrasquer, Luiza Oliveira, Gonzalo Hernandez, Mayka Sánchez, Fabien Delaspre, Elisa Cabiscol, Joaquim Ros, Jordi Tamarit

## Abstract

Friedreich Ataxia (FA) is a cardio-neurodegenerative disease caused by mutations in the frataxin gene, which result in low frataxin expression. It is well-established that frataxin deficiency affects iron homeostasis, but the tissue-specificity of these alterations is poorly understood. In this study, we have analyzed iron homeostasis alterations in the FXNI151F mouse model, which presents systemic frataxin deficiency and neurological defects resembling FA patients. Iron overload is observed in the brain from 21-week old FXNI151F mice, both males and females, and it does not further accumulate in older animals. It is also observed in livers from 39-week old mutant females, but not in males. Iron signaling is altered in all tissues: in brain and liver Iron Regulatory Protein 1 (IRP1) content is decreased, while in heart increased IRP2 and decreased specific aconitase 2 activity are observed. Remarkably, these cardiac alterations are partially restored in 39- week-old animals, suggesting that the heart is activating an iron-deficiency response to compensate for deficient iron-sulfur biogenesis. Our findings demonstrate that frataxin deficiency affects iron homeostasis in a time dependent and tissue-specific manner, and that the pathological mechanisms in FA comprise both iron accumulation and limited iron availability. Understanding this specificity is crucial for the design of iron-related therapeutic interventions aiming to improve FA symptomatology.

## INTRODUCTION

Friedreich Ataxia (FA) is an inherited recessive disease caused by mutations in the frataxin gene (*FXN*). The most common mutation is a GAA triplet expansion in the first intron of *FXN*, causing a systemic partial deficiency in frataxin protein expression. Around 4% of patients are compound heterozygous for a GAA expansion and a *FXN* point mutation or deletion. Clinical manifestations are mostly observed in the nervous system and the heart. Frataxin is mainly located in mitochondria, where it is involved in the biosynthesis of iron-sulfur proteins and/or in iron homeostasis. It is well established that its deficiency causes dysregulation of iron homeostasis, with iron overload first described in frataxin-deficient budding yeast (Babcock et al., 1997). Subsequent studies with this organism indicated that iron was accumulated due to an anomalous activation of the iron regulon (Lesuisse et al., 2003)(Moreno-Cermeno et al., 2010). Alterations in iron homeostasis have also been observed in patients and mammalian models of FA (Llorens et al., 2019; Ramirez et al., 2012; Whitnall et al., 2008). Regarding mice, iron aggregates were described in heart mitochondria from conditional cardiac/skeletal muscle frataxin KO mice (MCK mice) (Huang et al., 2009; Whitnall et al., 2012). These mice also presented changes in the expression of proteins involved in cellular iron homeostasis, such as transferrin receptor or ferritins, as well as Iron Regulatory Protein 2 (IRP2) activation. Iron Regulatory Proteins (IRPs) are activated upon iron deficiency and bind to Iron-regulatory Elements (IRE) present in the mRNAs from proteins involved in iron homeostasis. Such binding either protects mRNA from degradation (in the case of proteins involved in iron uptake such as Transferrin Receptor 1) or inhibits its translation (in the case of proteins involved in iron storage such as ferritins) (Sanchez et al., 2011). Elevated IRE-IRP2 binding activity was also observed in cultured fibroblasts derived from FA patients (Petit et al., 2021). Frataxin deficiency can also affect Iron Regulatory Protein 1 (IRP1), a protein that can be found in aconitase 1 (ACO1) or IRP1 forms, depending on the presence of an iron-sulfur cluster in its active site. ACO1 presents a [4Fe4S] cluster, which is required for aconitase activity, while IRP1 does not present the iron-sulfur cluster and adopts an open conformation that allows its binding to IRE elements. A marked deficiency in ACO1/IRP1 content was reported in liver-specific FXN KO mice (lvKO) (Martelli et al., 2015) and in the MCK mouse (Tong et al., 2022). The mechanisms causing alterations in IRPs in frataxin-deficient mammals are not completely understood. It is presumed that they are a consequence of deficient mitochondrial iron–sulfur biogenesis. Thus, prolonged loss of iron-sulfur clusters would result in decreased stability of ACO1/IRP1 (Tong et al., 2022), while IRP2 content would increase, as its degradation is mediated by the ubiquitin ligase FBXL5 which also contains an iron-sulfur cluster (H. Wang et al., 2020). Nevertheless, as iron-sulfur deficiency is not always observed in frataxin-deficient models, the involvement of other pathways in IRPs activation or destabilization should not be excluded.

As indicated above, alterations in iron homeostasis in mice have been mostly explored in conditional mouse models presenting null frataxin levels in the targeted tissues and normal expression in all the others. Although the results obtained have contributed to understanding the consequences of frataxin deficiency on iron metabolism, these conditional models do not fully replicate FA, since patients exhibit low frataxin levels across all tissues. We recently generated a mouse model based on the human pathological point mutation I154F (I151F in mice), which affects frataxin stability. Mice homozygous for this mutation (FXNI151F) present low frataxin levels in all tissues and display neurological and biochemical defects resembling those observed in FA patients (Medina-Carbonero et al., 2022; Sanz-Alcázar et al., 2023). Therefore, this model is an excellent tool to study the consequences of frataxin deficiency on different tissues. In the present work, we have used this model to explore the alterations in iron homeostasis in the brain (cerebrum and cerebellum), heart, and liver. We have found that all tissues present iron dyshomeostasis, but the precise alterations are tissue-specific.

## RESULTS

### Iron accumulates in the liver and the brain of FXNI151F mice

We had previously shown that in 21- and 39-week-old FXNI151F mice frataxin levels are between 3-6% of those observed in WT animals (Medina-Carbonero et al., 2022). By western blot we confirmed that at 10-weeks of age, similar values were observed, both in males and females (Sup. Fig. 1). To explore the consequences of frataxin deficiency on mouse iron homeostasis, we first analyzed total iron levels in two brain areas (cerebrum and cerebellum), heart and liver from WT and FXNI151F mice, at 10, 21 and 39 weeks of age. Tissues were digested in nitric acid and iron was quantitated by a bathophenanthroline-based assay. As shown in Figure 1, increased iron content was observed in the brain, while no significant changes were detected in the heart. Iron accumulation was observed earlier in the cerebellum than in the cerebrum, and it did not progress from 21 weeks onward. In the liver, marked differences in iron content were observed between males and females. Males presented a mild iron accumulation, which was significant in 21-week-old mice but did not progress in older mice (Sup. Fig. 2). Females presented a marked iron accumulation, which progressed in older animals (Fig 1 and Sup. Fig. 2). Increased liver iron content in females had been previously observed in wild-type mice (Krijt et al., 2004) and in murine models of hemochromatosis (Sproule et al., 2001), indicating that female mice are more prone to accumulate iron than males. In the brain and heart, these differences between males and females were not observed (supp Fig 1).

**Figure 1.**
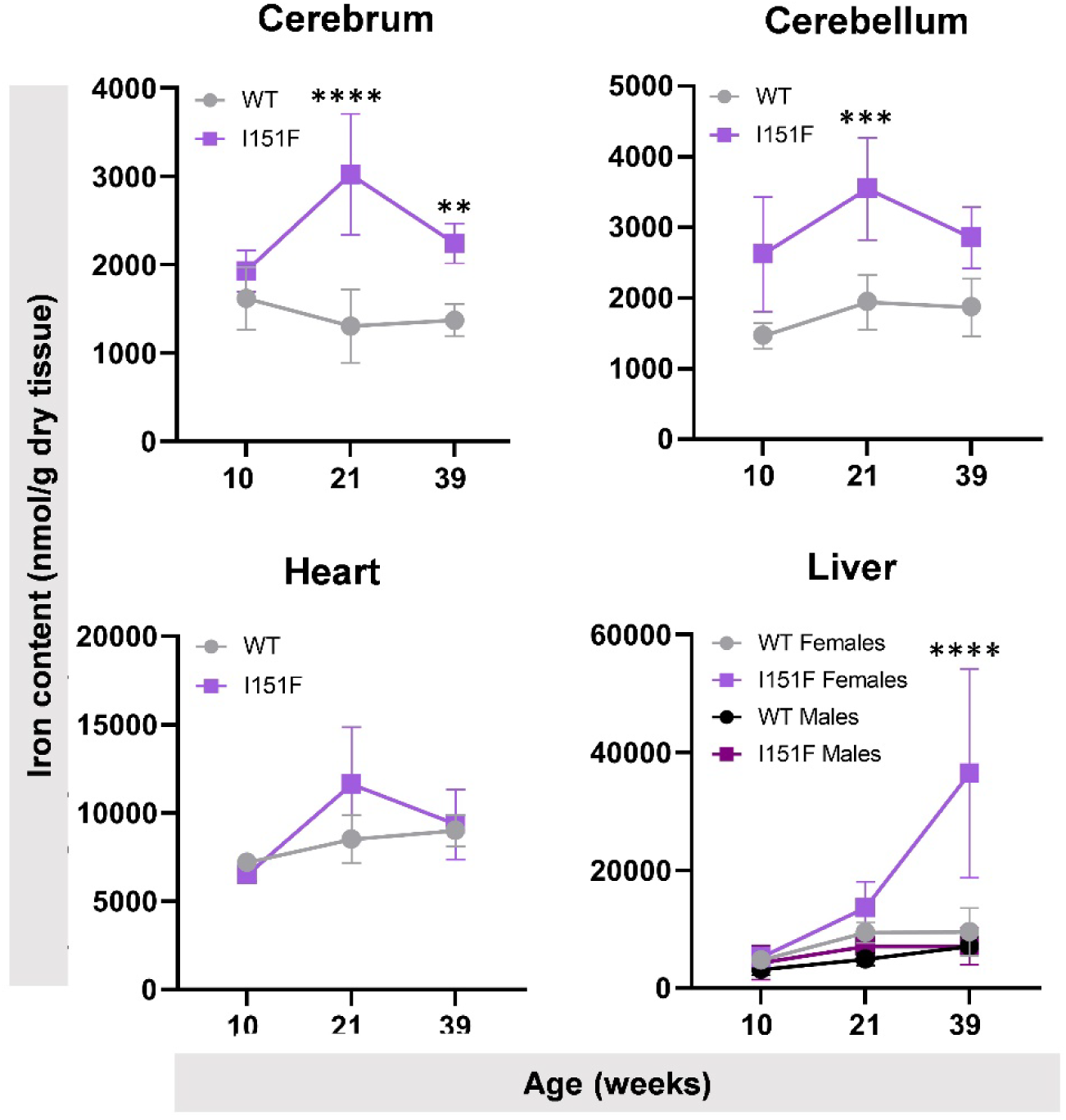
Non-heme iron levels in the cerebrum, cerebellum, heart and liver from WT and FXNI151F mice at indicated ages. For the liver, data of males and females are shown separately. Sex-separated data for the other tissues can be found in supplementary figure 2. Data are represented as mean +/- 95% confidence interval. For statistical analysis a two-way ANOVA test was done. Significant differences are indicated as p-values < 0.05(*), 0.01(**), 0.001(***) or 0.0001(****) between WT and FXNI151F mice.

### IRP1 and IRP2 content is altered in FXN I151F mice in a tissue-specific manner

To understand the mechanisms causing iron homeostasis disturbances, we analyzed Iron-regulatory proteins (IRP1 and IRP2). By western blot analyses, we observed decreased content of ACO1/IRP1 in the cerebellum, cerebrum and liver from 21-week-old FXNI151F mice, while no differences were observed in the heart (Fig 2A). In the liver, we evaluated if such decrease was similar between males and females, and we did not detect any difference in this parameter between sexes (Sup. Fig. 2). To test the activation status of IRP1 we performed an EMSA assay to assess the IRP1 binding capacity to a fluorescein-tagged IRE-containing mRNA probe. This assay was performed in liver homogenates, as the liver is the tissue with the highest expression of ACO1/IRP1. As a positive control, cellular extracts were treated with 2-mercaptoethanol (2ME) to fully activate IRP1. We could observe that the amount of active IRP1 (IRP1-IRE band) was decreased in the livers of FXNI151F mice, both in the presence or in the absence of 2ME (Fig. 2B). A lower amount of IRP1-IRE band in the positive control experiment (+ 2ME) confirmed that the total IRP1 amount was lower in mutant mice. We next evaluated IRP2 status. Interestingly, we observed increased content of IRP2 (thus, activation) in hearts from FXN I151F mice. In contrast, neither the liver, nor the brain presented altered IRP2 content (Fig. 2C). Therefore, we can conclude that IRP1 and IRP2 protein content is altered in FXNI151F mice in a tissue-specific manner: while IRP2 is activated in the heart, IRP1 is lower in the liver and the brain.

**Figure 2:**
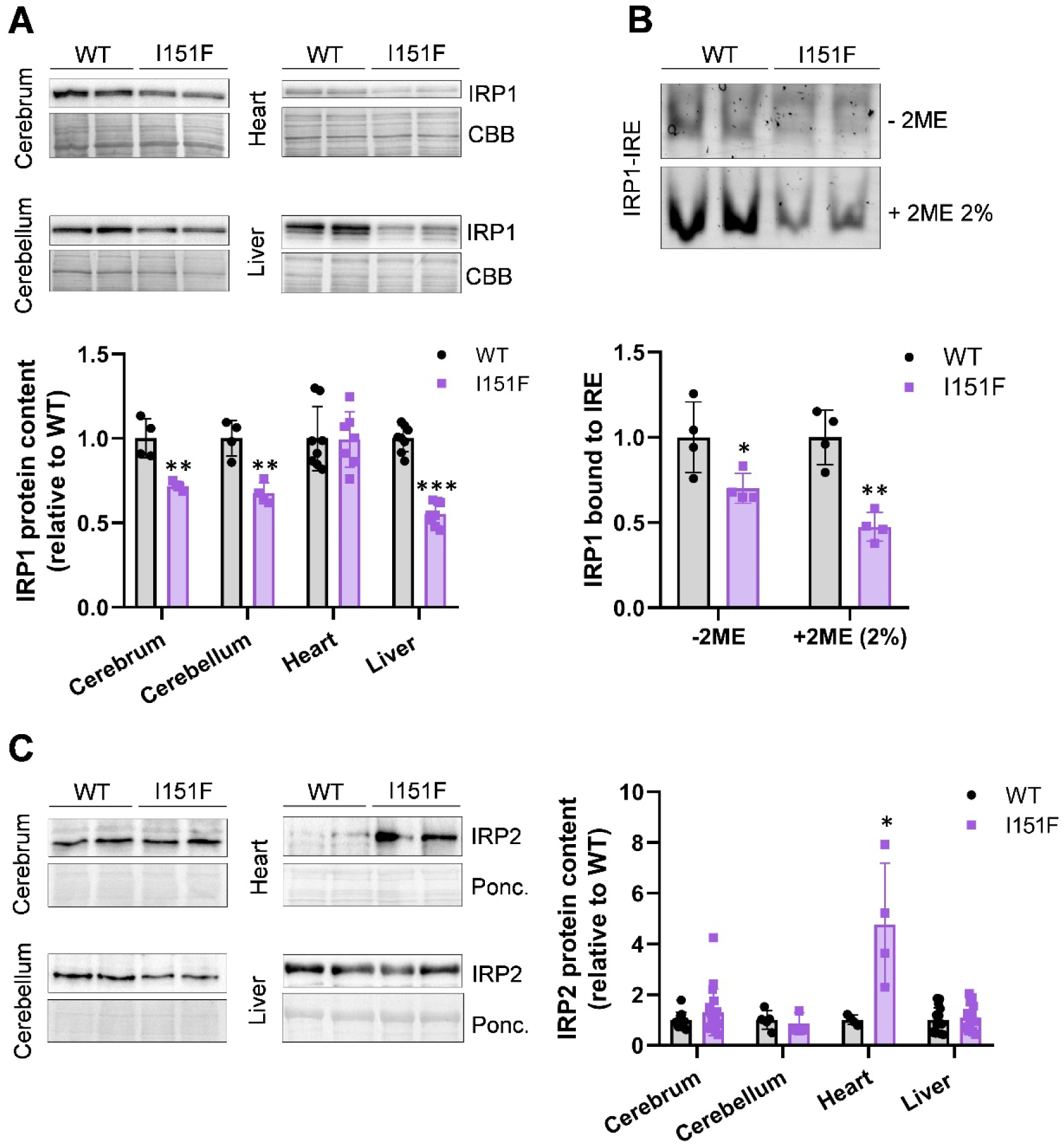
IRP protein status in 21-week-old mice. A) ACO1/IRP1 protein content was assessed by western blot in the indicated tissues. Post-western Coomassie blue staining (CBB) was used to assess protein load. Representative western blot images are shown above, while histograms represent the relative protein content of ACO1/IRP1, calculated from the western blot signal normalized to the CBB stain. B) EMSA analysis from liver samples. Representative assays showing the band corresponding to the IRE-bound IRP1 with or without including 2- mercaptoethanol (2ME). Histograms represent quantification (relative to WT values) of the IRP1- IRE band. C) IRP2 protein content was assessed by western blot in the indicated tissues. Representative western blot images are shown (left), while histograms represent the relative IRP2 protein content, calculated from the western blot signal normalized to the ponceau stain. All quantitative data are represented as mean +/- SD. Significant differences in p-values < 0.05(*) or 0.01(**), between WT and FXNI151F mice are indicated (t-test analysis).

### Analysis of IRP targets: Tfrc and ferritins

To investigate the consequences of dysregulation of IRPs on its targets, we measured TFR1 expression by qPCR (Gene: *Tfrc*), and heavy (FTH) and light (FTL) ferritins content by western blot, in the same tissues as above. *Tfrc* and *ferritins* mRNAs contain IREs, at 3’ (*Tfrc*) or 5’ (*ferritins*) from the coding region. Consequently, IRP binding protects *Tfrc* mRNA from degradation, while it inhibits the translation of *ferritins* mRNAs (Sanchez et al., 2011). We observed that *Tfrc* mRNA content was increased in FXNI151F mice hearts, while both ferritins protein content showed a tendency to decrease (Fig 3 B, C and D). These observations are in agreement with the induction of IRP2 observed in this tissue and suggest that an iron deficiency response is activated in the FXNI151F heart, despite the fact that total iron levels are not lower. A different response was observed in the liver and brain. Regarding the brain, the cerebrum did not show significant changes in either *Tfrc* or ferritins, while the cerebellum showed normal *Tfrc* expression and increased ferritins content. The liver presented increased *Tfrc* expression and ferritins content in both males and females (Sup Fig 2). These results confirm that alterations are tissue-specific. The increased Ferritins content in the cerebellum and liver could be due to lower IRP1 activity (described in the previous section). Nevertheless, the induction of *Tfrc* in liver is not compatible with the decreased IRP1 IRE-binding activity, suggesting alterations in other signaling pathways in the FXN151F mice apart from the IRP/IRE pathway.

**Figure 3:**
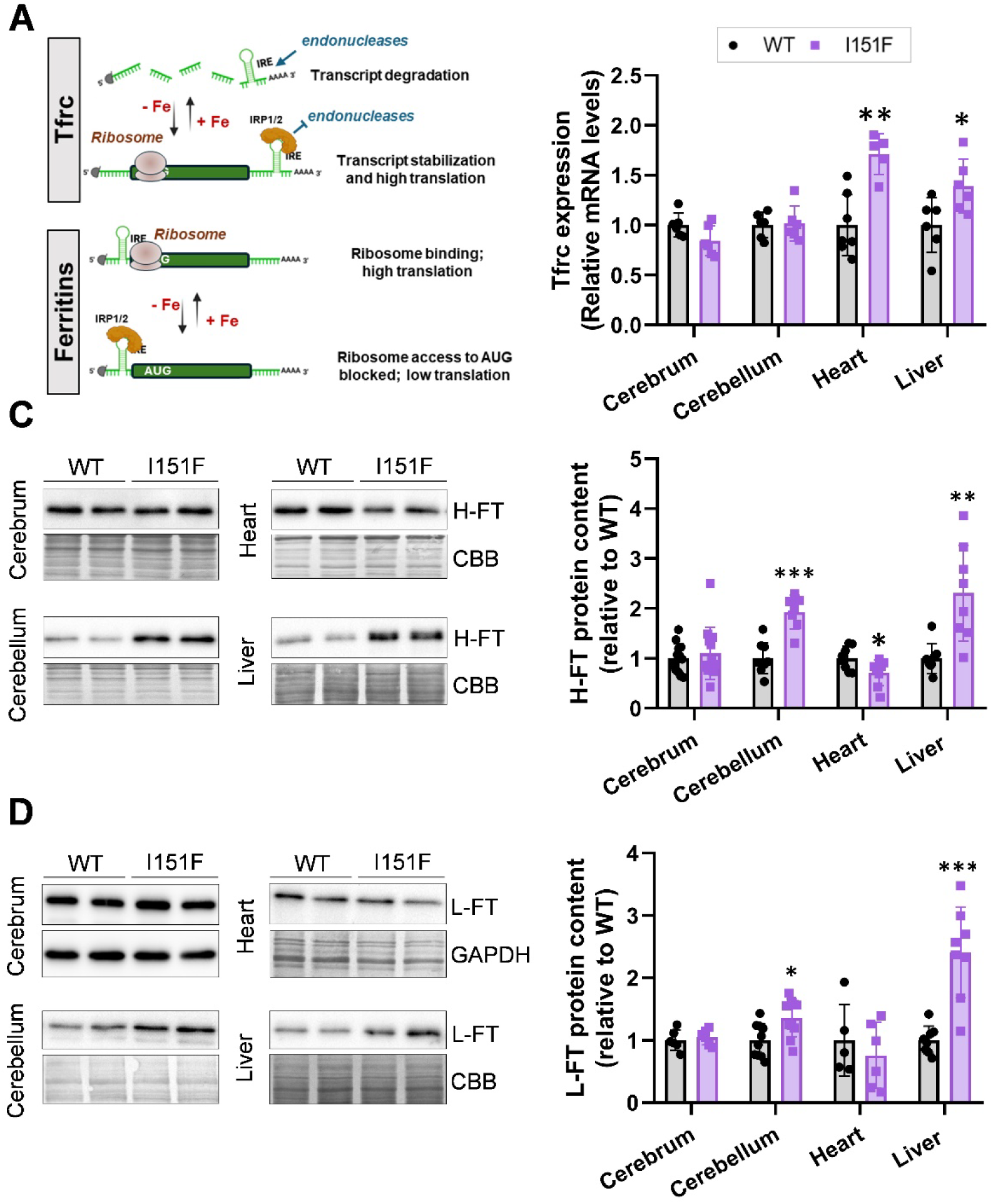
*IRP targets in 21-week-old mice. A) a simplified view of the IRP/IRE System. After iron deficiency, IRPs are activated and bind IRE elements, located at 3’ from the coding regions in Tfrc mRNA, or 5’ in Ferritins mRNA. This binding either blocks access to endonucleases that would cause transcript degradation (in the case of Tfrc) or impairs ribosome access to the start codon (in the case of ferritins). B) Tfrc mRNA content* was *measured by RT-qPCR. C and D) Ferritins heavy and light chains measured by western blot. Representative western blot images are shown in left panels, while histograms represent the relative protein content, calculated from the western blot signal normalized to the CBB stain or the GAPDH western blot signal. All quantitative data are represented as mean +/- SD. Significant differences p-values < 0.05(*), 0.01(**) or 0.001(***) between WT and FXNI151F mice are indicated (t-test analysis)*.

### Aconitase activities

We next decided to measure aconitase activities, commonly used as a marker of iron-sulfur status, and directly involved in the IRP/IRE pathways. Mammalian cells contain two aconitase isoenzymes: ACO1, which, as indicated before, is the iron-sulfur containing proteoform of IRP1 and is located in the cytosol, and aconitase 2 (ACO2), which is mitochondrial and its expression is regulated by the IRP/IRE pathway. According to data from the PaxDb database (M. Wang et al., 2015), ACO2 accounts for nearly 90% of total aconitase content in the brain and heart, while both isoenzymes are similarly expressed in the liver. To confirm these estimations, we first measured aconitase activity in the cerebellum, heart and liver from 21-week-old WT mice by an in-gel activity assay, which allows discrimination between both isoenzymes. Figure 4A shows the result obtained by this assay. The identity of both bands was verified by western blot from native gels run under the same conditions as the in-gel activity assays. It can be observed that ACO1 migrates faster than ACO2 and its activity is barely observed in the cerebellum and heart, which is consistent with the reported abundance data of these proteins in the PaxDb database. Consequently, the in-gel assay was only used to measure aconitase activity in the liver, while in the cerebellum and heart, we considered that ACO1 activity could not be reliably detected and that its contribution to total cellular aconitase activity was minimal. Therefore, for ACO2 activity estimation in these tissues, we decided to use a spectrophotometric assay that measures total aconitase activity (which, according to our in-gel assay is mostly contributed by ACO2 in the heart and cerebellum). This spectrophotometric assay is highly accurate, and allows the normalization of aconitase activity to citrate synthase (CS) activity, an indicator of total mitochondrial activity. The results obtained are shown in Figure 4. In the in-gel assay, a 50 % decrease in ACO1 activity was observed in the livers from mutant mice, while no significant decrease in ACO2 activity or protein content could be observed. Deficiency in ACO1 activity was more marked in females than in males (Fig. 4B). The spectrophotometric assay revealed a 25% decrease in total aconitase activity in this tissue (Fig 4C). This value is consistent with the 50% decrease in ACO1 activity and the preserved ACO2 activity found in the in-gel assay. Regarding heart and cerebellum, results from the spectrophotometric assay revealed a 27% decrease in ACO2/CS activity ratio in hearts from FXNI151F mice, while a 37% decrease was observed in cerebellum (Fig 4D). To determine if this decrease was caused by lower ACO2 protein content, or by decreased enzyme-specific activity, we measured the relative ACO2/CS protein content ratio by SRM-mass spectrometry, which provides accurate quantitation and allows the comparison of ratios between different proteins across multiple samples. Using this approach, we observed that the ACO2/CS protein content ratio was slightly increased in hearts from mutant mice, while a 36% decrease was observed in the cerebellum (Fig 4D). Therefore, specific ACO2 activity (calculated by dividing activity by protein ratios) was decreased in heart, but not in cerebellum. These results indicate that three different scenarios are observed regarding aconitase activities: in the cerebellum, decreased ACO2 activity is paralleled by decreased ACO2 protein content; in the liver, decreased ACO1 activity is paralleled by ACO1 protein loss, while ACO2 activity and content are not affected; finally, in heart decreased ACO2 activity is observed without loss of ACO2 protein content, indicating an inactivation of the enzyme. Such lower ACO2 specific activity in heart may indicate a decreased iron-sulfur biogenesis rate. Overall, these results confirm that the consequences of frataxin deficiency differ from one tissue to another.

**Figure 4:**
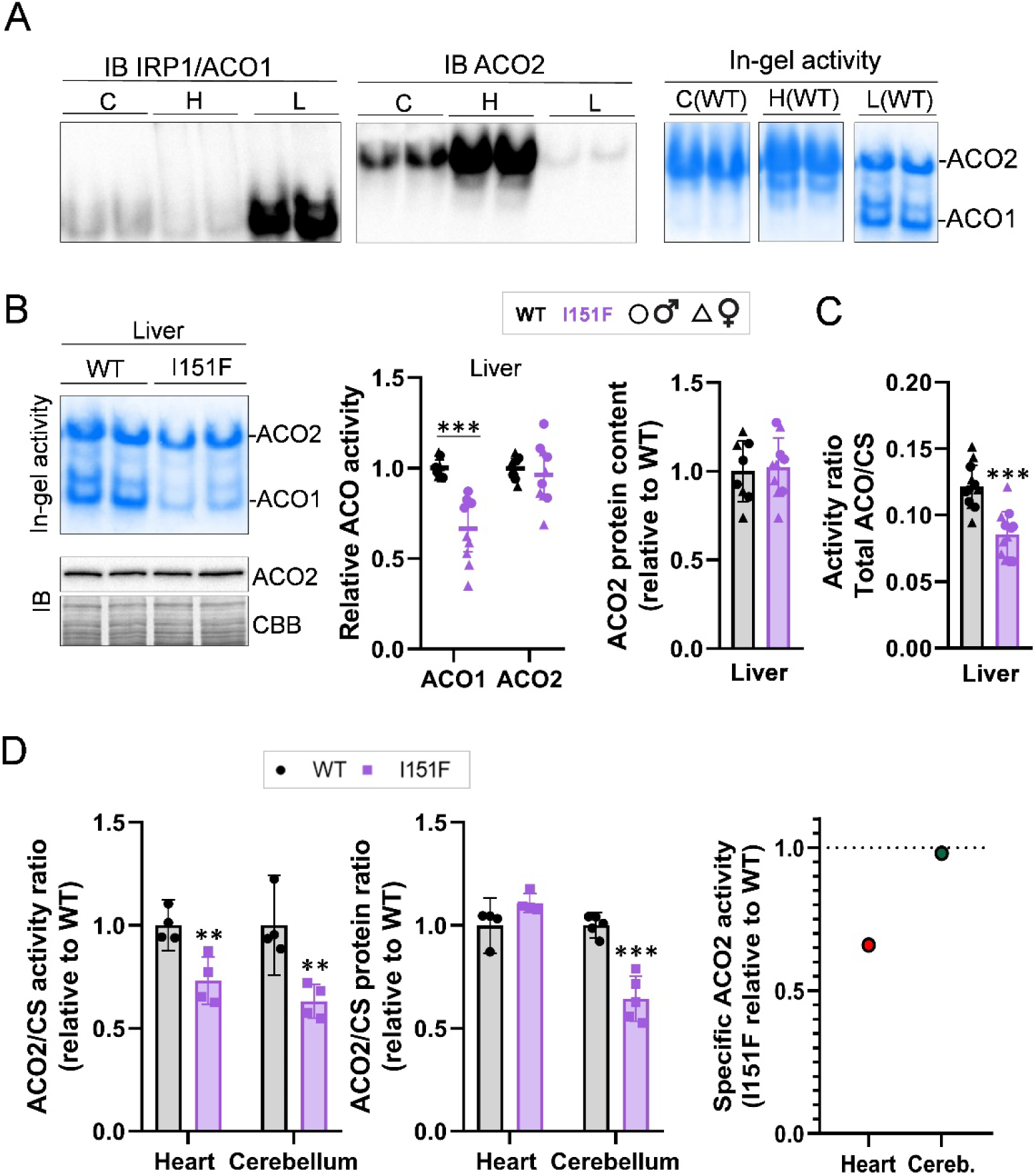
*Aconitase activities in 21-week-old mice. A) Migration of aconitase isoenzymes in the native gels used for in-gel aconitase activity. Homogenates from cerebellum (C), heart (H) and liver (L) obtained from WT mice (20* μ*g for heart and liver, 40* μ*g for cerebellum) were loaded on native gels and either assayed for aconitase activity or transferred to PVDF membranes for immunoblot (IB) analysis, which were incubated with antibodies against IRP1/ACO1 or against ACO2. Four bands can be detected in activity gels. Comparison with IB indicates that the two upper bands correspond to ACO2, while the two lower bands correspond to ACO1. This activity was barely detected in the cerebellum and heart, consistent with the low ACO1 IB signal observed in these tissues. B) Representative images from in-gel aconitase activity assay in the liver from WT and FXNI151F mice, and ACO2 western blot. Graphs represent the relative aconitase activity, calculated by the in-gel assays, and the ACO2 protein content, calculated from the western blot signal (normalized to the CBB stain). Individual male and female values are shown. C, ACO/CS activity ratio in the liver, as measured by the spectrophotometric assay. D, ACO2/CS activity and protein ratio in heart and cerebellum, as measured by the spectrophotometric assay (activity) and SRM-based mass spectrometry (protein). Specific ACO2 activity was calculated by dividing activity by protein ratios. All quantitative data are represented as mean +/- SD. Individual data points are indicated. In B and C, males are represented as circles, while triangles correspond to females. In D , both males and females are represented as squares Significant differences in p-values < 0.05(*), 0.01(**) or 0.001(***) between WT and FXNI151F mice are indicated*.

### ACO1/IRP1 is already decreased in 10-week-old mice

To uncover the mechanisms causing disturbances in ACO1/IRP1 and IRP2 in FXNI51F mice, we decided to analyze 10-week-old mice, in which iron accumulation is still undetected. In this regard, we hypothesized that young mice could provide information about the early events causing alterations in iron homeostasis. In these mice we found no changes in the content of IRP2 in any of the tissues analyzed, indicating that IRP2 induction in the 21-week-old heart is a later event (Fig. 5A). Regarding ACO1/IRP1 protein content, we observed that it was already decreased in liver and cerebellum, while in the cerebellum such decrease was less marked at 10 weeks compared to 21-week-old mice (Fig. 5A). Moreover, cerebrum ACO1/IRP1 was not decreased, indicating that the loss of this protein in the brain is progressive, and that cerebellum is more affected than cerebrum. We decided to explore the liver in more detail, as it was the tissue with a more marked loss in ACO1/IRP1. EMSA analysis revealed decreased content of active and total IRP1, confirming that ACO1/IRP1 content was lower, and revealing that the remaining IRP1 was not activated in FXNI151F mice (Fig. 5B). Nevertheless, when we analyzed ferritins protein content and *Tfrc* mRNA levels in this tissue, we observed a slight increase in *Tfrc* expression, and decreased content from both ferritins (Fig. 5C).

**Figure 5:**
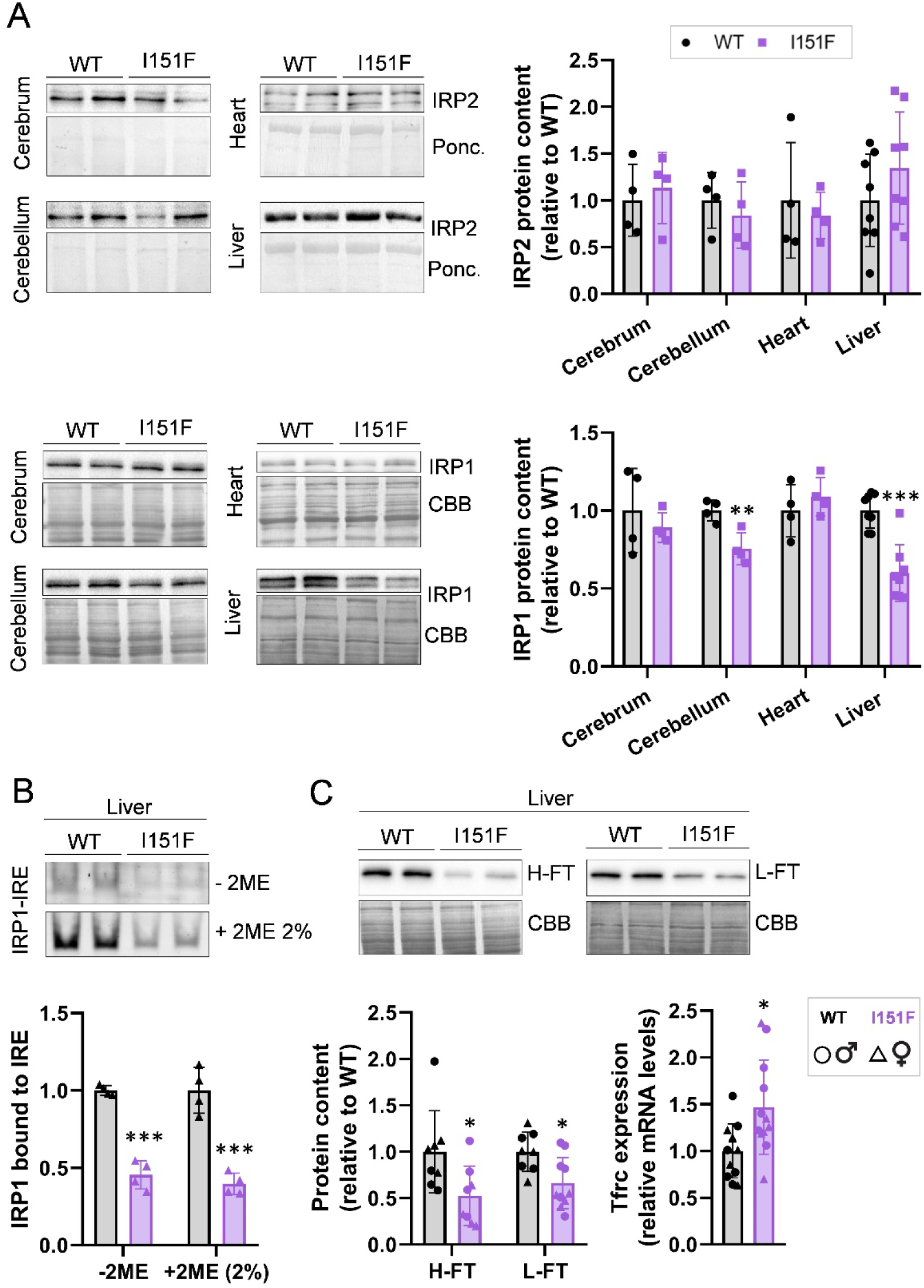
IRP status in 10-week-old mice. A) IRP2 and ACO1/IRP1 protein content was assessed by western blot in the indicated tissues. Representative western blot images, as well as quantification are shown. B) EMSA analysis from liver samples. Representative assays showing the band corresponding to the IRE-bound IRP1 in the presence and absence of 2ME and quantification of the IRP1-IRE band are shown. C) IRP targets in liver: Ferritins’ protein content was assessed by western blot, while Tfrc expression was measured by qPCR. All data are represented as mean +/- SD. Individual data points are indicated. In A, both males and females are represented as squares. In B and C, males are represented as circles, while triangles correspond to females.

This signature is typical from an iron deficiency response. However, this response cannot be attributed to IRPs, as none of them was found activated in 10-week-old mice, confirming that there may be alterations in additional signaling pathways.

### Iron-sulfur protein status in 10-week-old liver

ACO1/IRP1 deficiency has been previously described in several FA models, and attributed to decreased iron-sulfur biogenesis. Nevertheless, the presence of active ACO2 protein in the 21-week-old liver suggests that in conditions of partial frataxin deficiency, the cause of ACO1/IRP1 protein loss may not be an impairment in mitochondrial iron-sulfur biogenesis. To obtain a more comprehensive picture of the iron-sulfur status in the liver, we evaluated the content of several iron-sulfur proteins in 10- week-old mice. We measured aconitase activity by the in-gel assay, and we observed decreased ACO1 activity and normal ACO2 activity, as in 21-week-old mice (Fig. 6A). ACO2 protein content was not affected, as revealed by western blot (Fig. 6C and D). We also analyzed other iron-sulfur-containing proteins from different cellular compartments, including the NDUFS1 complex I subunit, the SDHB complex II subunit and ferredoxin 1 (FDX1), located in mitochondria, Glutaredoxin-3 (GLRX3), amidophosphoribosyltransferase (PPAT) and dihydropyrimidine dehydrogenase (DPYD), which are cytosolic, and the catalytic subunit from DNA polymerase delta 1 (POLD1), located in the nucleus. As mitochondrial controls, we measured the UQCRC2 subunit from mitochondrial complex III and the ATPA subunit from complex V. We observed decreased content of DPYD in FXNI151F mice and normal levels of the other proteins analyzed (Fig. 6C and D). These results confirm that partial frataxin deficiency does not cause a general loss in iron-sulfur proteins. Instead, it affects specifically some of them in a tissue-specific manner. Therefore, we can conclude that ACO1/IRP1 decreased protein content in the liver is not caused by a general loss in iron-sulfur clusters.

**Figure 6:**
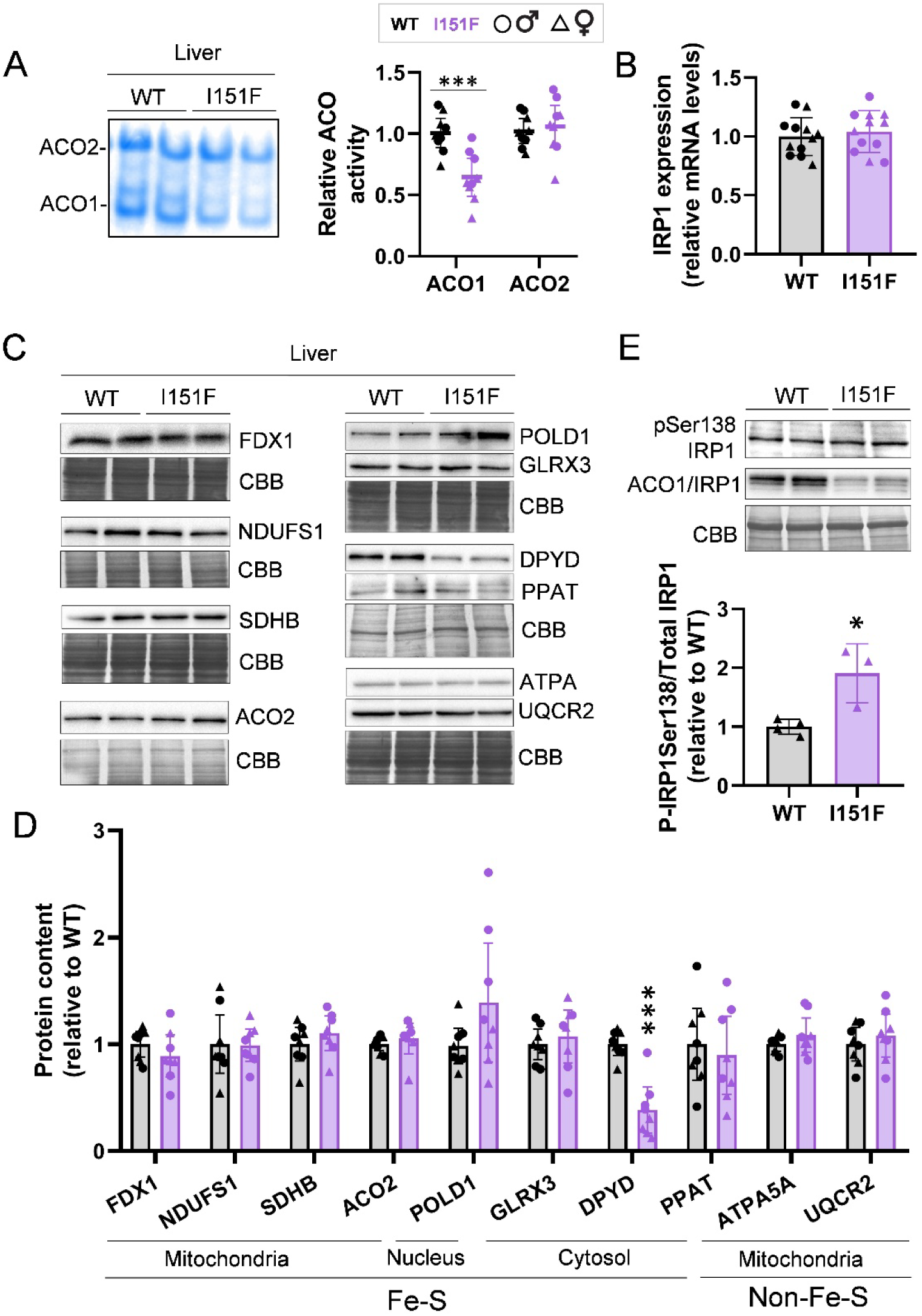
Iron-sulfur protein status in 10-week-old mice livers. A) ACO1 and ACO2 activities were measured in native gels. Representative images and quantifications are shown. B) ACO1/IRP1 mRNA expression was measured by qPCR. C and D) content from the indicated proteins was assessed by western blot. Representative western blot images are shown in C, while quantification is shown in D. E, representative western blot images show the detection of total ACO1/IRP1 and a phosphorylated proteoform (pS138IRP1) by specific antibodies. The histogram represents the relative quantification of the pIRP1/total IRP1 ratio. All data are represented as mean +/- SD. Individual data points are indicated: circles correspond to females, while triangles correspond to males.

### ACO1/IRP1 is phosphorylated in liver from 10-week-old FXNI151F mice

As stated above, results from the EMSA assay and the presence of remaining ACO1 activity (indicated by the in-gel assay), suggest that ACO1/IRP1 deficiency may not be caused by a general iron-sulfur loss. To further explore the causes of ACO1/IRP1 loss in the liver, we measured its mRNA expression by qPCR, where we did not observe any decline, indicating that protein loss is caused by post-transcriptional events (Fig. 6B). As phosphorylation of this protein at Ser-138, has been linked to increased sensitivity to degradation (Fillebeen et al., 2003), we decided to explore the ACO1/IRP1 phosphorylation status using antibodies specific for phospho-Ser138. As shown in Figure 6E, the intensity of the phospho-Ser138 band was similar between WT and FXNI151F mice. As total ACO1/IRP1 is decreased, we can conclude that increased phosphorylation of this residue is occurring in the livers of FXNI151F mice. Therefore, our results suggest that frataxin deficiency in the liver alters signaling pathways, which trigger ACO1/IRP1 phosphorylation and cause increased sensitivity to degradation of this protein.

### IRP2 induction recovers iron-sulfur deficiency in the heart

As shown in Fig 4D, we detected decreased specific ACO2 activity in hearts from 21- week-old mice, suggesting that iron-sulfur cluster biogenesis is compromised in this tissue. Such decreased iron-sulfur biogenesis could be caused by lowered cysteine desulfurase activity (an enzyme required for iron-sulfur biogenesis) as its activity can be stimulated by frataxin *in vitro* (Patraa & Barondeaua, 2019)). Nevertheless, induction of IRP2 and *Tfrc* was also observed, indicating that heart was activating an iron-deficient response. These observations suggest that the decline in specific ACO2 activity could also be caused by limited iron availability. To clarify this issue, we decided to analyze the changes in IRP2 content, *Tfrc* expression and specific ACO2 activity at different ages. Therefore, we analyzed ACO2/CS activity/protein ratios, *Tfrc* expression and IRP2 content in heart at 10 and 39 weeks of age, and compared them with those obtained in 21-week old mice. Both ACO2/CS activity and protein ratios presented a slight (≈ 5 to 15%) decrease in hearts from 10-week and 39-week old mutant mice, while specific ACO2 activity was not affected at 10-weeks and presented a ≈10% decrease at 39 weeks (Fig. 7A). Moreover, IRP2 content and *Tfrc* expression were partially restored to WT levels in 39-week old I151F mice (Fig 7B and C). The evolution of these parameters from 10 to 39-week-old animals is summarized in Fig 7D. It can be observed that IRP2 and *Tfrc* follow a similar pattern, were maximum increase is observed in 21-week old mutant animals. Interestingly, specific ACO2 activity present an inverse pattern, with maximal decrease at 21 weeks, and partial recovery at 39 weeks of age. These observations support the hypothesis that ACO2 deficiency in 21-week old heart is caused by limited iron availability, which would activate an iron deficiency response that by increasing iron uptake would provide iron for synthesizing iron-sulfur clusters and recovering ACO2 specific activity.

**Figure 7:**
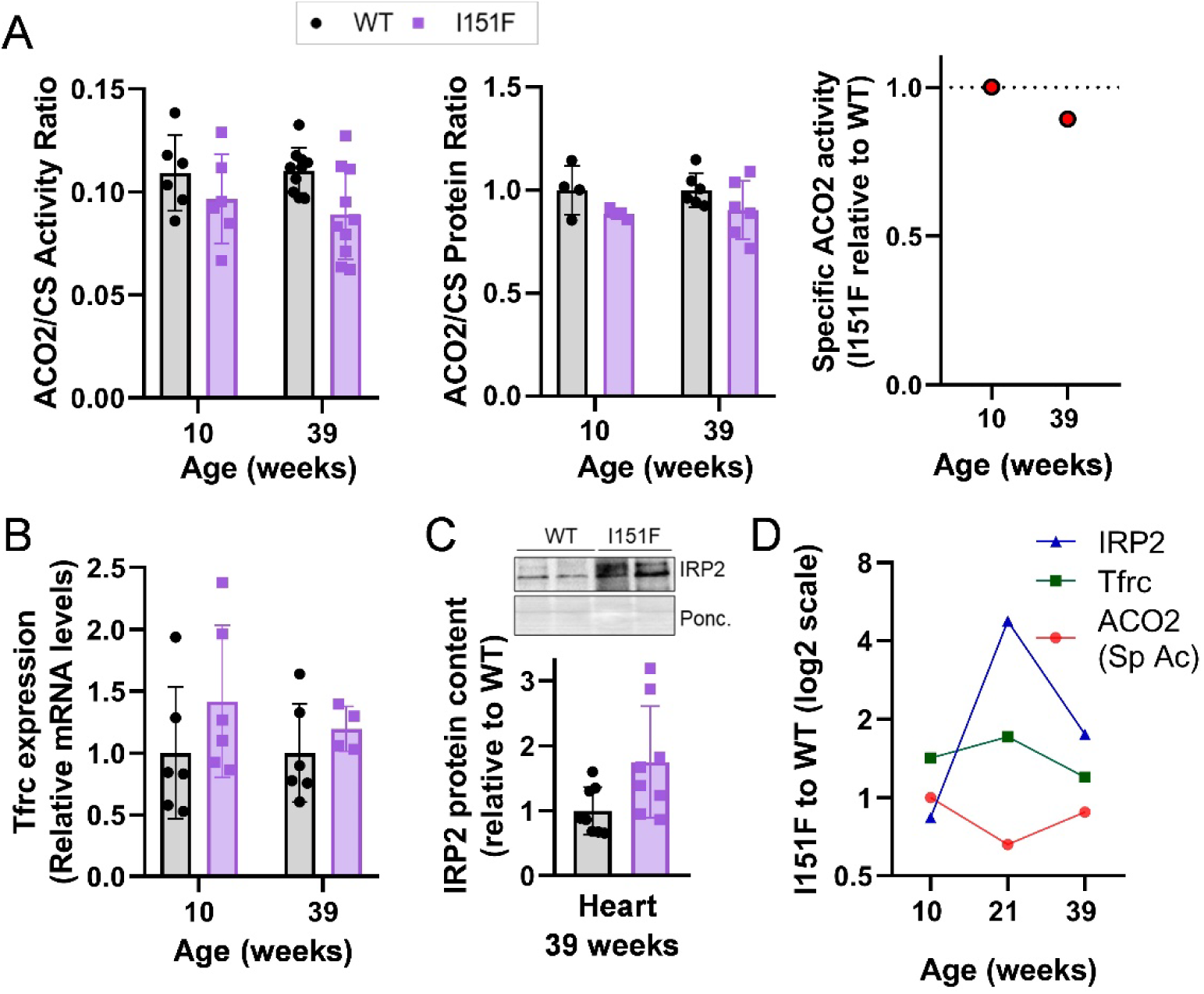
Evolution of IRP2 content and aconitase activity in the heart. A) ACO2/CS activity and protein ratio, as well as ACO specific activity, measured by a spectrophotometric activity assay and by SRM-proteomics. B) Tfrc expression in 10 and 39-week-old hearts analysed by qPCR. C) IRP2 content in 39-week-old heart, detected by western blot. D) Evolution of relative values (I151F to WT) of IRP2 content, Tfrc expression and specific ACO2 activity from 10-week to 39- week- old mice.

## DISCUSSION

The consequences of frataxin deficiency on iron metabolism had previously been analysed in heart or liver-specific KO frataxin mice. However, these conditional models did not accurately replicate the human pathological condition, as patients present partial frataxin deficiency in all the tissues. In the present work, using the new FXNI151F murine model, we have analysed for the first time the disturbances to mammalian iron homeostasis caused by systemic partial frataxin deficiency, thus approaching the conditions experienced by FA patients. We have mainly focused on the cerebellum, heart and liver. The cerebellum and heart are among the most affected tissues in FA, while the liver has a crucial role in systemic iron homeostasis. Some analyses have also been performed in the cerebrum, a different brain area, to compare its affectation with the one experienced by the cerebellum.

The results obtained indicate that alterations in iron homeostasis are time-dependent and tissue-specific, with early iron accumulation in the cerebellum and a later accumulation in the liver of female mice. In the heart, where total iron content is not altered, IRP2 content and *Tfrc* expression are increased. The induction of both proteins had previously been described in the hearts of cardiac KO mice (MCK mice), which presented iron overload and decreased ferritins and ACO2 content (Whitnall et al., 2012). In this regard, we have observed partial ACO2 activity deficiency in this tissue, without loss of ACO2 protein content. Overall, our findings resemble those obtained in the MCK mouse, despite the observed changes being less intense in our model. This difference can be because our model has a partial frataxin deficiency while the MCK model has a total deficiency. In this sense, our results would be more representative of the situation experienced by patients and highlight that under conditions of partial frataxin deficiency the heart shows an iron deficiency response. The observation that under conditions of partial frataxin deficiency, the reduction in ACO2 activity is not due to a loss of protein but possibly to a loss of its iron-sulfur centre is particularly interesting. Such loss may be consequence of impaired iron-sulfur biogenesis due to lowered cysteine desulfurase activity (required for iron-sulfur biogenesis and whose activity is stimulated by frataxin *in vitro* (Patraa & Barondeaua, 2019)) or of limited iron availability due to the formation of iron complexes or aggregates (which would render this metal unavailable for the iron-sulfur biogenesis machinery, causing a paradoxical iron deficiency condition). Such anomalous iron complexes or aggregates have been observed in hearts of the MCK mice (Whitnall et al., 2012) and in frataxin-deficient yeast (Seguin et al., 2011). Uncovering the precise mechanism causing ACO2 deficiency in 21-week-old hearts is beyond the objectives of the present work. Nevertheless, the absence of ACO2 deficiency in 10-week-old mice and its recovery in 39-week-old mice support the hypothesis of limited iron availability for iron-sulfur biogenesis, as decreased cysteine desulfurase activity should have a similar impact on ACO2 protein at younger and older ages. Besides, the comparison between 10-week, 21-week and 39-week-old mice highlights the relationship between ACO2 deficiency, IRP2 activation, and *Tfrc* expression. It further supports the hypothesis of limited iron availability in frataxin-deficient hearts, which would partially be alleviated by increasing iron uptake through *Tfrc* induction.

In this work, we have also observed decreased ACO1/IRP1 protein content in the liver and the brain. Such loss is tissue-specific, as it does not occur in the heart and is progressive, as it is much accentuated in 21-week than in 10-week-old brains. Loss of this protein had previously been observed in other FA models. In a model of inducible frataxin deficiency in a human embryonic kidney cell line, it was observed that ACO1/IRP1 decreased earlier than mitochondrial ACO2 (Lu & Cortopassi, 2007), highlighting the relationship between frataxin and ACO1/IRP1. More recent studies have reported ACO1/IRP1 deficiency in both the liver (Martelli et al., 2015) and the cardiac/skeletal muscle (Tong et al., 2022) frataxin conditional KO mouse models. Since ACO1/IRP1 devoid of a cluster can be ubiquitinated and degraded (J. Wang et al., 2007), it was assumed that such deficiency would be a consequence of impaired iron-sulfur biogenesis. However, this hypothesis does not fit with the data obtained in our model, as the remaining ACO1/IRP1 protein is active as an aconitase (as evidenced by in-gel assays), and IRE-binding activity does not increase in livers from 10-week-old FXNI151F mice. Moreover, ACO2 and other iron-sulfur proteins are not affected in liver, as would be expected if a general deficiency in these cofactors was the cause of ACO1/IRP1 loss in this tissue. These observations suggest that the degradation of this protein may be caused by a mechanism independent of iron-sulfur cluster deficiency. Interestingly, studies in budding and fission yeasts indicate that iron–sulfur deficiency in frataxin-deficient yeasts is amplified upon activation of the iron regulon, which triggers a metabolic reprogramming program that downregulates iron-containing proteins (Alsina et al., 2018; Gabrielli et al., 2012; Moreno-Cermeño et al., 2013). Thus, ACO1/IRP1 deficiency in brain and liver could also be caused by the alteration of regulatory pathways. Indeed, the phosphorylation at ser138-IRP1 points to that direction. Moreover, decreased ACO1/IRP1 protein content has been reported in liver biopsies from hereditary hemochromatosis patients (Neonaki et al., 2002), in macrophages treated with paraquat (Milczarek et al., 2017) , and in livers of SOD1-deficient mice (Starzyński et al., 2005). In this last work It was suggested that down-regulation of ACO1/IRP1 could be caused by phosphorylation of ACO1/IRP1 at Ser-138 by protein kinase C, which would be activated by oxidative stress. ACO1/IRP1 levels can also be regulated by the MEK/ERK and PI3K/Akt pathways (Zhang et al., 2014). All these observations highlight the relevance of phosphorylation events in ACO1/IRP1 regulation and indicate that ACO1/IRP1 deficiency can be observed in conditions where either iron homeostasis or oxidative stress are altered. Dysregulation of signaling events upon frataxin deficiency could explain the specificity observed regarding tissue, age, or iron-sulfur protein inactivated, as signaling networks show high specificity. In other words, if ACO1/IRP1 loss was exclusively caused by impaired iron-sulfur biogenesis, this phenomenon would probably be observed in all conditions analysed, irrespective of age or tissue, and would affect most iron-sulfur proteins.

In addition to ACO2 and ACO1/IRP1, we have analyzed other iron-sulfur-containing proteins in the liver, including Glrx3, a multifunctional iron-sulfur cluster binding protein whose function is related to cytosolic iron-sulfur cluster assembly and delivery (Banci et al., 2015). It has been reported that its depletion decreases the content and/or activity of several extramitochondrial iron-sulfur proteins, including IRP1 (Cheng et al., 2023). However, we have only observed DPYD deficiency, confirming the specificity of frataxin deficiency consequences. Nevertheless, this is an interesting observation, as mutations in this protein can cause dihydropyrimidine dehydrogenase deficiency (DPYDD), a condition that shows large phenotypic variability, ranging from no symptoms to a convulsive disorder with motor and mental retardation. In addition, both homozygous and heterozygous mutation carriers can develop severe toxicity after the administration of the antineoplastic drug 5-fluorouracil (5FU), which is catabolized by the DPYD enzyme (Van Kuilenburg et al., 1999). Therefore, the possibility of a partial DPYD deficiency in FA patients should be taken into account when considering antineoplastic pharmacological treatment administration. Besides, as urinary excretion of uracil and thymine is increased in DPYDD patients, it would be interesting to test these compounds in urine from FA patients for use as biomarkers in clinical trials.

Despite being well-established that frataxin deficiency disturbs iron homeostasis, remains unclear the specific role of iron in the pathogenesis of FA.. Iron-mediated toxicity has been reported in several models of the disease (Tamarit et al., 2021). However, symptoms or markers commonly associated with iron deficiency are also found in FA patients, such as serum ferritin values in the lower range or restless leg syndrome (Grander et al., 2024), a clinical condition associated with iron depletion (Frauscher et al., 2011). Moreover, IRP1 activation (which promotes iron uptake) has a protective effect in a mouse model of hepatic frataxin deficiency (Martelli et al., 2015), while dietary iron supplementation ameliorates cardiac hypertrophy in the cardiac conditional frataxin KO mutant mice (Whitnall et al., 2012). Iron chelation therapies have provided mild beneficial effects at low doses, but a worsening of the condition at higher doses of the drug (Elincx- Benizri et al., 2016; Pandolfo & Hausmann, 2013). Overall, these observations suggest that the pathological mechanisms in FA could be related to both iron accumulation and limited iron availability. In this context, the results obtained in the present work indicate that systemic frataxin deficiency in mice affects iron homeostasis in a tissue-specific way. This specificity could be behind the contradictory observations regarding the role of iron in FA. Our results provide some clues to characterize this specificity which may contribute to the design of therapeutic interventions that aim to improve FA symptomatology.

## EXPERIMENTAL PROCEDURES

### Animals

FXNI151F heterozygous mice (C57BL/6J-Fxnem10(T146T,I151F)Lutzy/J) were obtained from the Jackson Laboratory, Bar Harbor, ME, USA (Stock Number 31922) as previously described (Medina-Carbonero et al., 2022). Intercrosses of heterozygous animals were performed to generate the homozygous WT and FXNI151F mice. Animals were housed in standard ventilated cages with 12 h light/dark cycles and fed with a regular chow diet ad libitum. Genotyping was performed by sequencing the *Fxn* gene PCR product amplified from DNA extracted from tail biopsy specimens as previously described (Medina-Carbonero et al., 2022).

### Tissue Iron quantification

Tissue non-heme iron was measured using the iron chelator BPS (bathophenanthroline disulfonic acid) as previously described (Tamarit et al., 2006) with some modifications. A tissue fragment of approximately 30 mg was dried in a SpeedVac at 60°C and weighed. To this sample, 212 μl of MilliQ water and glass beads (diameter between 0.5 and 1 mm, Sigma G8772) were added, and tissue was homogenized using a BioSpec Mini-Beadbeater. Then, 288 μl of 70% nitric acid was added (resulting in a final concentration of nitric acid at 40%), and the mixture was incubated for 2 h at 98°C in a Thermomixer comfort (Eppendorf) before a 5 min centrifugation at 10,000 rpm. The supernatant (60 μl) was diluted with 740μl of MilliQ water. After, 400μL of this dilution were mixed with 160 μl of 38 mg/ml sodium ascorbate, and 126 μl of ammonium acetate (saturated ammonium acetate diluted 1/3 with water). Volume up to 1mL was completed with water. The absorbance of this solution at 535 nm and 680 nm was read before and after adding 15 μl 34 mg/ml BPS, and the increase in absorbance caused by the formation of the BPS-iron complex was used to calculate the iron content per dry weight (nmols Fe per mg of dry tissue). Blanks and standards (prepared with ferrous ammonium sulfate) were also analyzed to prepare standard curves and correct for potential iron contamination in the materials and reagents used.

### Western blot

Between 20-100mg of tissue were minced into 2–3 mm^2^ pieces and, placed in 1.5 ml screw cap polypropylene tubes in the presence of lysis buffer consisting of 50 mM tris(hydroxymethyl)aminomethane (Tris) HCl pH 7.5, protease inhibitor cocktail and PhosphoSTOP (Roche). 375 μl of lysis buffer per 100mg of tissue was used. The mixture was homogenized in a BioSpec Mini-Beadbeater with Glass beads (0.5–1.0 mm) After, SDS was added to the mixture at 4% final concentration. This homogenate was vortexed for 1 minute, heated at 98°C for 5 minutes, sonicated and centrifuged at 14000rpm for 10 min. Protein content in the supernatant was quantified using the Pierce^TM^ BCA Protein Assay Kit (ThermoFisher Scientific) according to the manufacturer’s instructions. After SDS-polyacrylamide gel electrophoresis, proteins were transferred to PVDF (Millipore, IPVH00010) or Nitrocellulose (Sigma_Aldrich, 10600093) membranes and blocked with 3% I-block (ThermoFisher, T2015) for at least 1 hour. Membranes were probed with the following primary antibodies overnight: Frataxin 1:1000 (AbCam, ab219414), IRP1/ACO1 1:10000 (AbCam ab183721), IRP2 1:1000 (Novus Biologicals, NB100-1798), H-FT 1:1000 (AbCam, ab75973), L-FT 1:10000 (abCam, ab69090), ACO2 (Sigma, HPA001097), FDX1 1:1000 (AbCam, ab108257), NDUFS1 1:60000 (AbCam, ab108257), OxPhos Rodent Antibody Cocktail 1:3000/1:40000 (Invitrogen, 458099) for SDHB, UQCR2 and ATPA5A detection, POLD1 1:10000 (AbCam, AB186407-1001), GLRX3 1:1000 (Proteintech, 11254-1-AP), DPYD 1:1000 (Santa Cruz, sc-376681), and PPAT 1:1000 (LifeTechnologies, PA5-27770). Proteins were detected after incubating with peroxidase conjugated secondary antibodies for 1 hour and 5min with Immobilon^R^ Western (Millipore) after 5x washes. Images were acquired by ChemiDoc MP system from Bio-Rad. Membranes were stained with Coomassie brilliant blue (CBB) or Ponceau for normalization. When required, data was analysed using ImageLab software (Bio-Rad).

### ACO2/CS protein ratio quantification by SRM-proteomics

Tissue homogenates obtained as described above (100 ug of protein) were precipitated with cold acetone (9 volumes) and resuspended in 1% sodium deoxycholate, 50 mM ammonium bicarbonate. Then, proteins were subjected to reduction by 5,3 mM DTT and alquilation by 26 mM iodoacetamide. Proteins were digested overnight at 37°C with mass spectrometry grade trypsin (SOLu-Trypsin, Sigma) in an enzyme:substrate ratio of 1:50. After, formic acid was added to precipitate sodium deoxycholate. The resulting peptide mix was purified and enriched using 100ul Pierce C18 ZipTips. Eluted fraction from the C18ZipTip was evaporated using a Concentrator Plus (Eppendorf) and peptides were resuspended in 3% acetonitrile plus 0,1% formic acid containing heavy peptide standards from ACO2 (NAVTQEFGPVPDTAR and DLEDLQILIK) and CS (GLVYETSVLDPDEGIR and DYIWNTLNSGR) obtained from JPT (SpikeTidesTM_L). All peptide samples were analyzed on a triple quadrupole spectrometer (Agilent 6420) equipped with an electrospray ion source. Chromatographic separations of peptides were performed on an Agilent 1200 LC system using a Supelco Bioshell A160 Peptide C18 column (1 mm x 15 cm). Peptides (up to 15 micrograms of protein digest) were separated with a linear gradient of acetonitrile/water, containing 0.1% formic acid, at a flow rate of 50 μl/min. A gradient from 3 to 60% acetonitrile in 45 minutes was used. The mass spectrometer was operated in multiple reaction monitoring mode. Transitions were obtained either from peptide atlas, SRM atlas or Prosit (Gessulat et al., 2019) imported into Skyline software (MacLean et al., 2010) and validated with the heavy internal standards. Skyline was also used to analyze results. For calculating the ACO2/CS protein abundance ratio, the light to heavy (L/H) ratio obtained for each peptide in each replica was divided by the mean average L/H value of each peptide among all samples and replicas to obtain a normalized L/H value. The normalized L/H value from the different peptides corresponding to the same protein and sample was averaged, and finally sample ACO2/CS ratio was calculated.

### Enzyme activities

Spectrophotometric analysis of Aconitase/citrate synthase activity ratios was performed as previously reported (Medina-Carbonero et al., 2022). Briefly, tissues were minced into 2–3 mm^2^ pieces and placed into tubes containing non-denaturing lysis buffer consisting of 50mM Tris-HCl at pH 7.4, protease inhibitor cocktail (Roche) and 2,5mM sodium citrate. A BioSpec Mini-Beadbeater was used for tissue homogenization with Glass beads (0.5–1.0 mm). Then, Triton X-100 was added at a final concentration of 0.5%. Homogenized tissues were centrifuged for 5 min at 13000 rpm at 4°C and the supernatants were obtained. Aconitase activity was measured in 50mM Tris-HCl at pH 7.4, containing 1 mM of sodium citrate, 0.2 mM of NADP, 0.6 mM of manganese chloride and 0.25 units of isocitrate dehydrogenase (Sigma-Aldrich, I2002). NADPH formation was measured at 340 nm for 120s. Citrate synthase activity was measured with a coupled assay to reduce 5,5′-dithiobis-(2-nitrobenzoic acid) (DTNB). Briefly, tissue extracts were added to Tris-HCl 100 mM pH 8.1 with 0.4 mg/ml of DTNB and 10 mg/ml of Acetyl-CoA. Absorbance was measured at 412 nm during 120 s. Then, 8.5 mg/ml of oxaloacetate was added into the cuvette and the absorbance was measured again at 412 nm for 120 s for the detection of reduced DTNB. Values are presented as a ratio of aconitase activity versus citrate synthase activity. In-gel aconitase activity analysis was performed as described before (Ghosh et al., 2008) with some modifications. Tissue homogenates were prepared as above (in this case with 50mM Tris-HCl pH 8.5). 20 μg of total protein was loaded on a gel composed of a separating gel containing 6% acrylamide, 132 mM Tris base, 66 mM borate, 3.6 mM citrate, and a stacking gel containing 4% acrylamide, 66 mM Tris base, 33 mM borate, and 3.6 mM citrate. The running buffer contained 25 mM Tris (pH 8.3), 96 mM glycine, and 3.6 mM citrate. Electrophoresis was carried out at 170 V at 4°C for 2h 45min. After, gels were incubated for 10-15 min at 37°C, dark conditions, in 10 ml of a 100mM Tris-HCl pH 8 solution containing 1mM NADP, 2,5mM cis-aconitic acid, 5mM manganese chloride, 0,3mM phenazine methosulfate and 2,6 U/ml of isocitrate dehydrogenase until blue bands were observed. Images were acquired in a GelDoc EZ imager (Bio Rad) and band intensity measured by ImageLab software. Relative aconitase activity was quantified by comparison with a standard curve prepared by loading different amounts of WT homogenates on gels.

### Quantitative real-time PCR

For qRT-PCR analysis, 50-100 µg of tissue samples were homogenized with 1 mL of TRIzolTM Reagent (Invitrogen, 15596018) using an IKA® T10 basic homogenizer. RNA was extracted following the manufacturer’s instructions. For each sample, 1 µg of total RNA was converted into cDNA with iScript cDNA Synthesis Kit (Bio-Rad, 1708891) and 50 ng of cDNA was used in each reaction. Assays were performed in a CFX96 Real-Time System (Bio-Rad) using TaqMan® 2X Universal PCR Master Mix (Applied Biosystems, 4304437) mixed with TaqMan probes: Tfrc1 (Mm00441941_m1), Aco1 (Mm00801417_m1) and GAPDH (Mm99999915_g1) or Rpl19 (Mm02601633_g1) as a loading control. Quantification was completed using Bio-Rad CFX Manager real-rime detection system software (version 3.1, Bio-Rad). Relative expression ratios were calculated based on ΔCp values with efficiency correction considering multiple samples.

### EMSA assay

Cytosolic extracts were isolated from liver (1-2 mm^3^ pieces) in 0,5mL of ice-cold cytoplasmatic lysis buffer (CLB) buffer (25 mM Tris-HCl pH=7,4, 40mM potassium chloride, 1% Triton x100, 1mM DTT and Complete EDTA free protease inhibitor cocktail (Roche), with a Dounce tissue homogenizer for 10 s. After chilling on ice for 20min and centrifuging for 10min full speed at 4°C, supernatants corresponding to cytosolic fractions were obtained. Then, the protein was quantified by Bio-Rad protein assay (ref. 5000006) and 25μg of total protein was mixed with CLB up to 10μl and 1μM FAM-labeled FTH-1 IRE probe previously heated for 1 min at 95°C. The FAM-labeled fluorescent FTH-1 IRE probe with the sequence 5’- UCCUGCUUCAACAGUGCUUGGACGGAAC-3’ was prepared by GenScript Biotech (Netherlands). β-mercaptoethanol was added before the probe to the corresponding positive controls for total IRP binding activity. After 20min incubation, a 10-minute incubation with 1 μl of 50mg/ml heparin was done, both at room temperature in dark conditions. Components were separated using 5% TBE gels with a 1:60 acrylamide: bis ratio, at 100V for 45 min. Subsequently, image acquisition was performed in a ChemiDoc MP system from Bio-Rad with blue light epi illumination and a 530/28 emission filter.

### Statistical analyses

For statistical analyses, GraphPad Prism (version 8) was used. Differences between groups in iron content were assessed by two-way ANOVA with the Tukey’s multiple comparison test when considering more than two categorical variables into the analyses. Two-tailed Student’s t test was used to assess the significance of differences between WT and I151F mice in proteins, mRNA expression, EMSA and aconitase activity experiments. Significant p values are indicated as <0,05(*), <0,01(**), <0,001(***), <0,0001(****).

## Authors’ contributions

MPG and MMC performed most of the experiments. ASA, MPC FD and LO assisted in the collection and preparation of mouse tissues. EC, GH and MS assisted and supervised in the biochemical analyses. All authors provided technical support and suggestions for the project and for the manuscript. JR and JT conceived the project and supervised the study. MPG, MMC and JT designed the experiments, analyzed and interpreted data, and wrote the manuscript.

## Acknowledgements

We thank Roser Pané for technical assistance, as well as the Proteomics and the Animal services from Universitat de Lleida, and the metabolomic services from IRBLLeida.

## Abbreviations

FA: Friedreich Ataxia
IRP: Iron regulatory protein
FT: Ferritin
ACO: Aconitase
CS: Citrate Synthase
EMSA: Electrophoretic mobility shift assay
CBB: Coomassie brilliant blue

