## Supplemental Figures for "SYSTEMIC FRATAXIN DEFICIENCY CAUSES TISSUE-DEPENDENT IRON HOMEOSTASIS ALTERATIONS: IMPLICATIONS FOR FRIEDREICH ATAXIA"

#### Supplemental Fig 1

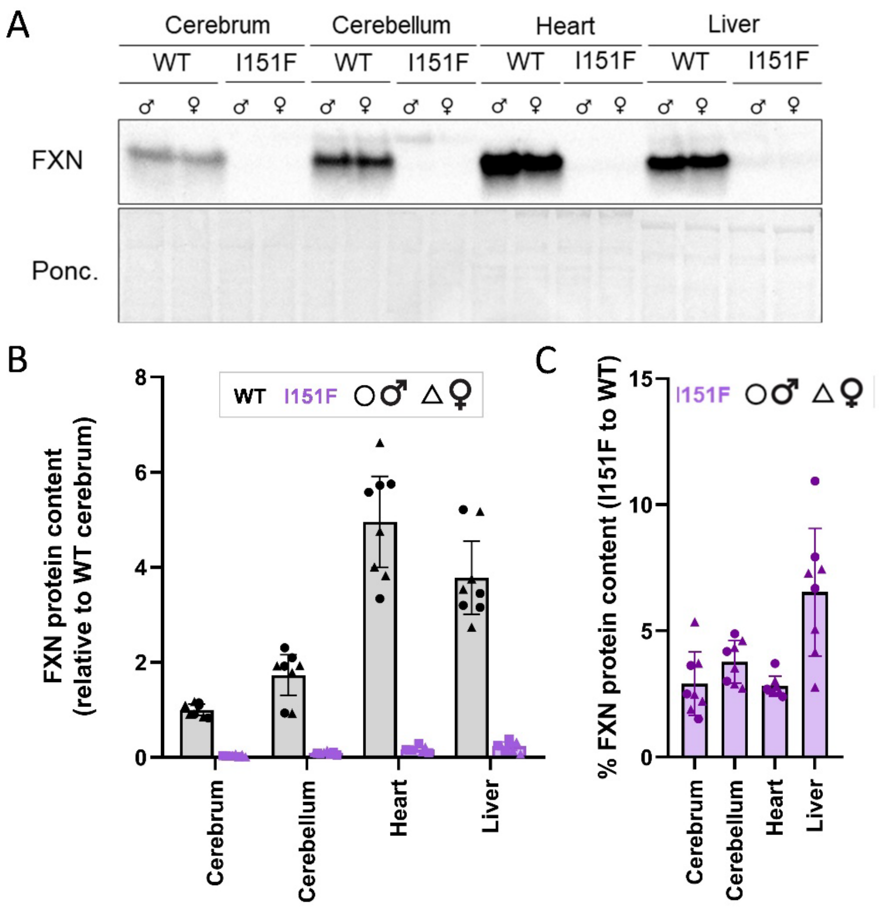


***Supplemental figure 1. Mature frataxin content analyzed by western blot in homogenates from 10-week old WT and FXN$^{I151F}$ mice.*** *A) Representative frataxin western blot image showing detection of mature frataxin in homogenates from cerebrum, cerebellum, heart and liver. Samples from two WT and two FXN$^{I151F}$ mice per group (one male and one female) were loaded on the same gel. B) Frataxin content relative to the values obtained in WT cerebrum (mean±SD) calculated from at least 8 independent measurements. C) Remaining frataxin content in the indicated tissues from I151F mice, as compared to WT.*

Supplemental Fig 2

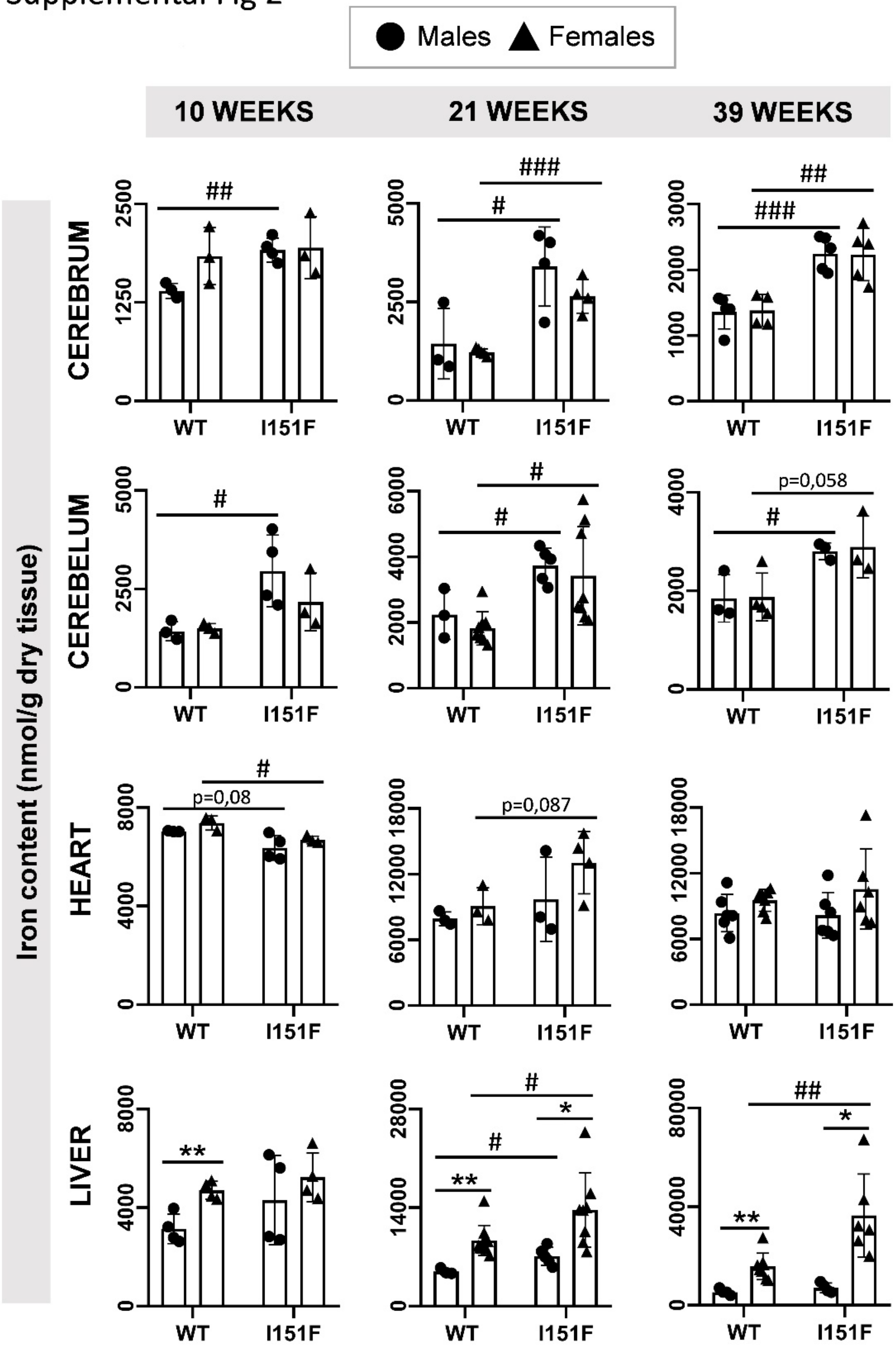


*Supplemental figure 2- Differences in iron content between males and females. Non-heme iron was quantified in the indicated tissues from mice at different ages. Data is represented as mean +/- SD. Significant differences between females and males are indicated by * (p < 0.05) or ** p < 0.01. Significant differences between WT and I151F are indicated by # (p < 0.05), ## (p < 0.01) or ### p < 0.001.*

Supplemental Fig 3

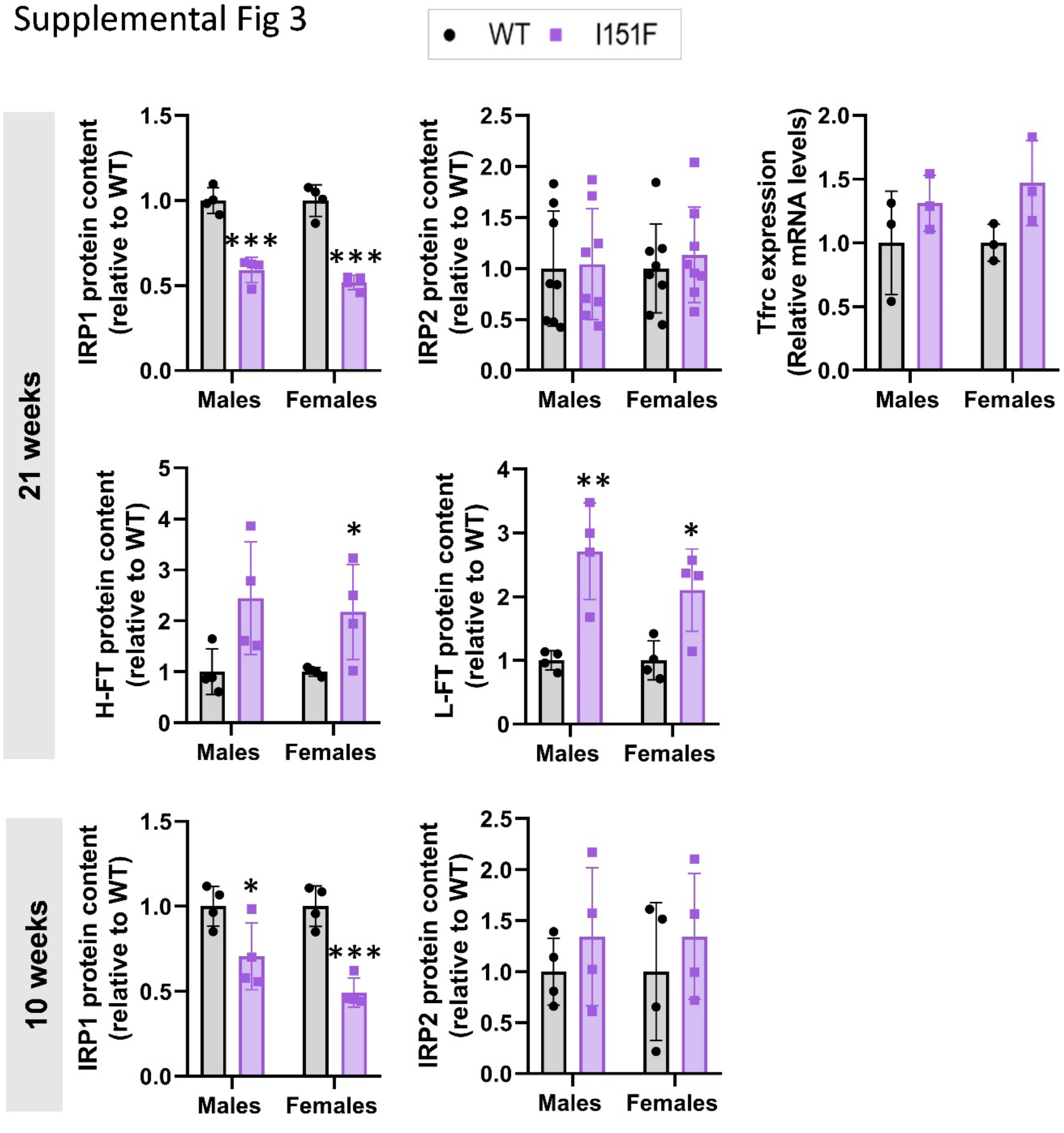


*Supplemental Figure 3 – Impact of sex on several parameters analyzed on liver. Values are those shown in figures 2, 3 and 5 without differentiating males and females. Significant differences between WT and I151F mice are indicated. No significant differences between males and females were observed in any of the parameters analyzed.*

### Supplemental Fig. 4

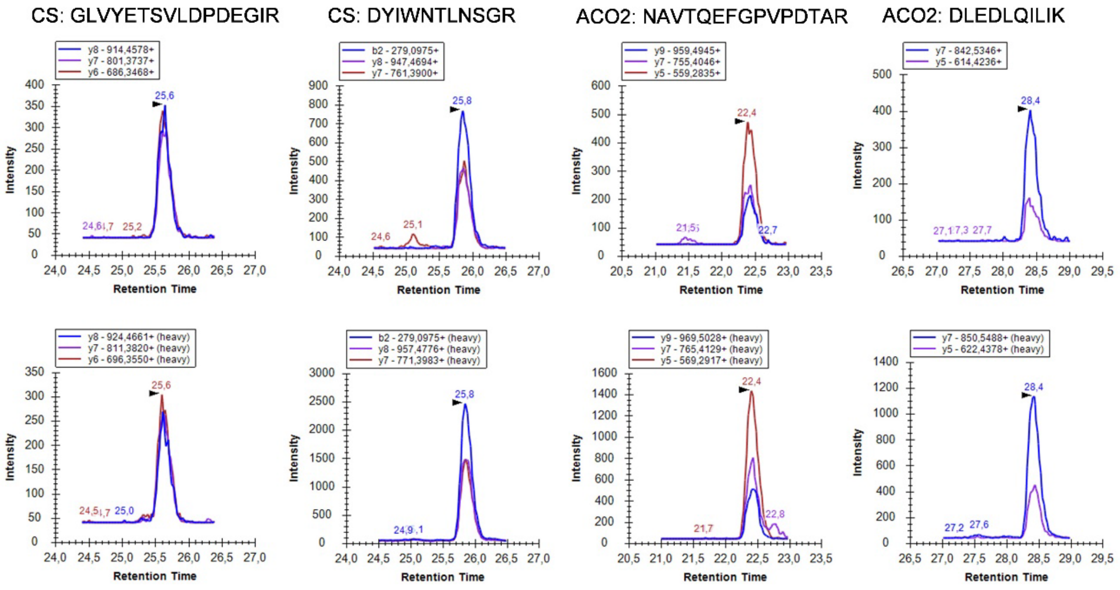


*Supplemental fig. 4. Representative SRM traces used to quantify ACO2/CS protein ratio. Two peptides from citrate synthase (CS) and Aconitase 2 (ACO2) were analyzed. The upper trace corresponds to the endogenous peptide (light version), while the lower trace corresponds to the internal standard (heavy version). Images correspond to peptides obtained from a 21 week old WT mice heart sample.*
